# Structural basis of nick translation in human DNA replication

**DOI:** 10.64898/2026.07.22.739974

**Authors:** Inas Zein, Ammar U. Danazumi, Muhammad Tehseen, Lingyun Zhao, Kerry Blair, Rania Almaghrabi, Vlad-Stefan Raducanu, Samir M. Hamdan, Alfredo De Biasio

**Author notes:** These authors contributed equally (Inas Zein, Ammar U. Danazumi, Muhammad Tehseen).

## Abstract

Nick translation during Okazaki fragment maturation requires iterative coordination of DNA polymerase delta (Pol δ), which displaces the downstream primer, and flap endonuclease FEN1, which cleaves the resulting flap, on the sliding clamp PCNA. The structural basis of this coordination is unknown. We present cryo-EM structures of human Pol δ–PCNA, Pol δ–PCNA–FEN1 and FEN1–PCNA on flap DNA, capturing four states of the nick translation cycle. Pol δ strand displacement emerges from structural elements intrinsic to the B-family fold rather than dedicated separation machinery, with a conserved palm loop acting as separation wedge and PCNA engagement required for melting of the downstream duplex. In the Pol δ–PCNA–FEN1 toolbelt, FEN1 is pre-positioned opposite Pol δ on PCNA to receive the substrate. Nucleotide removal triggers DNA handoff while both enzymes remain clamp-bound, followed by Pol δ dissociation. A post-handoff structure reveals stable DNA retention by FEN1–PCNA after flap cleavage, explaining the slow nick translation kinetics and the obligate role of Ligase 1 in sealing.

## Introduction

Lagging-strand DNA replication is inherently discontinuous: because DNA polymerases synthesise exclusively in the 5′-to-3′ direction, the lagging strand is replicated as a series of short (∼200 nt) Okazaki fragments that must subsequently be joined into a continuous daughter strand^1^. In eukaryotes, this maturation process is carried out by a dedicated set of enzymes. Each fragment is initiated by DNA primase–polymerase α (Pol α), which synthesises a short RNA–DNA primer of ∼20–30 nucleotides on the single-stranded template exposed by the replicative helicase ^2–4^. These primers are extended by the replicative polymerase Pol δ ^5^, which associates with the homotrimeric sliding clamp PCNA to achieve high processivity^6,7^. Upon encountering the 5′ end of the downstream RNA–DNA primer, Pol δ transitions from gap-filling synthesis to strand displacement, executing nick translation — an iterative process in which Pol δ displaces a short stretch of the downstream primer while simultaneously extending the upstream strand, generating a 5′ flap of 1–2 nucleotides that is cleaved by flap endonuclease 1 (FEN1) to regenerate a nick^8,9^. Following each cleavage event, Pol δ resumes synthesis to displace the next nucleotide stretch, and the cycle repeats until the RNA–DNA primer is fully removed. The resulting nick is then sealed by DNA Ligase I (Lig1)^8,10,11^. The iterative nature of nick translation requires coordination between Pol δ and FEN1 on PCNA.

Despite extensive biochemical characterisation, the structural basis for several key steps of nick translation remains unresolved. First, how Pol δ achieves strand displacement is unclear. Pol δ belongs to the B-family of DNA polymerases, shares no structural homology with the separation elements described in A-family polymerases ^12,13^, and its displacement activity is weak, clamp-dependent, and prone to stalling at the nick junction^8,9,14^. Second, how FEN1 is spatially and temporally coordinated with Pol δ on PCNA to cleave short flaps before they anneal or become extended is unknown. Third, whether substrate handoff from Pol δ to FEN1 occurs via a toolbelt mechanism — in which both enzymes simultaneously occupy PCNA — or via sequential exchange has been debated^8,9,15^. Fourth, biochemical data indicate that nick translation by human Pol δ is slow and that Lig1 is required to displace the nicked product from FEN1 for sealing^8^ yet the structural basis of this rate-limiting step is missing.

A previous cryo-EM structure of the human Pol δ–PCNA–FEN1 complex assembled on primer– template DNA^16^ resolved the architecture of the toolbelt but did not capture the strand-displacement-competent state, the exchange-competent intermediate, or a post-handoff product.

Here we present four cryo-EM structures of human Pol δ–PCNA, Pol δ–PCNA–FEN1, and FEN1–PCNA complexes assembled on flap DNA substrates that mimic discrete stages of the nick translation cycle. Together, these structures provide a detailed view of how strand displacement is achieved by a B-family polymerase, how Pol δ and FEN1 are coordinated on PCNA, and how the post-cleavage state of FEN1 sets the stage for Lig1-mediated ligation.

## Results

### Structural basis of strand displacement by Pol ***δ***–PCNA

To determine the structural basis of strand displacement by the Pol δ holoenzyme, we assembled Pol δ– PCNA on a flap DNA substrate with a chain terminator in the upstream primer strand, purified the complex by gel filtration, and added the next correct nucleotide (dTTP) immediately prior to cryo-EM grid preparation. Cryo-EM reconstruction yielded a 3.1 Å map resolving Pol δ, the PCNA protomer bound to Pol δ, and the upstream and downstream duplex segments (Figure 1a and Extended Data Figure 1; Extended Data Table 1), with residual DNA and PCNA heterogeneity recovered by 3DVA^17^ (Figure 1a). Similarly to the ternary complex assembled with primer–template DNA ^16^, the catalytic subunit of Pol δ (p125) rests atop PCNA, with the regulatory subunits (p50, p66, and p12) arranged laterally and DNA extending from the active site through the clamp ring (Figure 1b). PCNA heterogeneity stems from rigid-body rotations of the ring around the flexible polymerase–clamp binding interface (Figure 1b), as reported previously for the holoenzyme bound to primer-template DNA ^16^.

**Figure 1.**
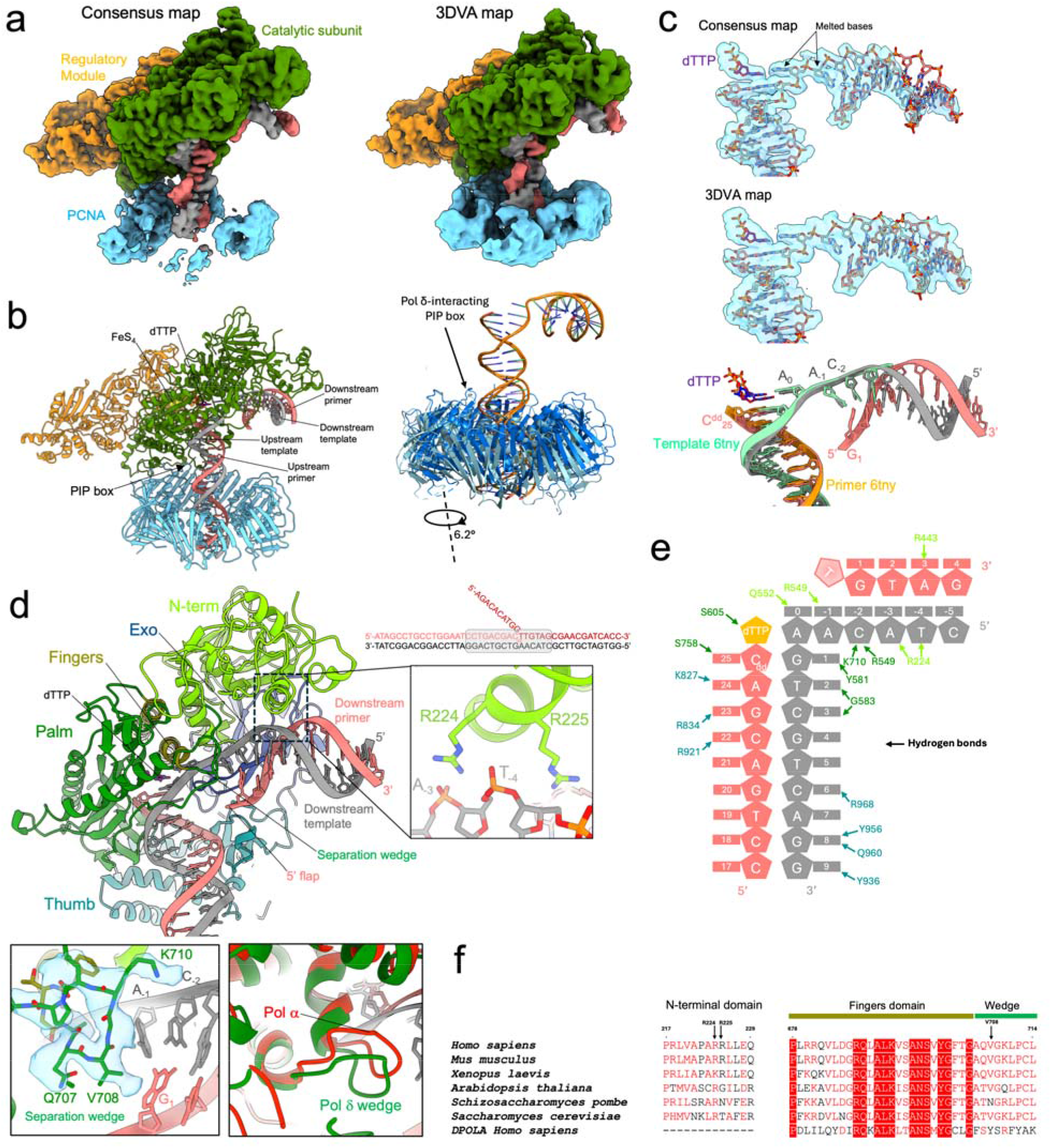
Structural basis of flap engagement by Pol δ–PCNA. **(a)** Cryo-EM consensus and 3DVA map of the Pol δ holoenzyme bound to flap DNA and dTTP. **(b)** Molecular model of the complex, shown to the left of an overlay of the PCNA and DNA models for the two extreme 3DVA frames from the first component, highlighting rigid-body rotation of the PCNA ring. **(c)** Cryo-EM consensus and 3DVA map of the junction region of the flap substrate. The inset shows an overlay with the primer–template DNA substrate from the Pol δ complex^16^ (PDB: 6TNY), indicating that engagement of the flap substrate results in melting of two downstream base pairs. **(d)** Model of the Pol δ catalytic domain engaging the flap substrate, with subdomains labelled. The inset to the right highlights basic residues of the N-terminal domain forming a positively charged pocket adjacent to the template strand of the downstream duplex. The sequences of the oligonucleotides used in the substrate are shown above, with the region interacting with the polymerase shaded in grey. The inset to the bottom left shows an enlarged view of the map and model of the separation wedge and nearby region. The inset to the bottom right shows an overlay of Pol δ and Pol α catalytic domains, highlighting structural conservation of the separation wedge. **(e)** Schematic representation of the interactions between Pol δ and the flap substrate, with polymerase residues coloured according to subdomain. **(f)** Sequence alignment in the N-terminal and fingers/wedge regions of Pol δ homologues and human Pol α.

The primer terminus adopts a canonical engaged configuration with bound dTTP, closely matching the ternary complex assembled on primer–template DNA ^16^ (Figure 1c). The downstream duplex exits the palm domain at an approximately right angle and is positioned adjacent to a positively charged pocket in an α-helix of the N-terminal domain, formed by R224 and R225 (Figure 1c-e); R224 is conserved across Pol δ homologues (Figure 1f) and establishes direct contacts with bases A-3 and T-4 of the template strand (Figure 1d-e). The 10 nt 5’ ssDNA flap projects outward from the polymerase and is conformationally flexible and therefore not resolved (Figure 1d). Pairing of dTTP with its complementary template base destabilises the corresponding nucleotide on the displaced strand (Figure 1c-e), and the adjacent downstream base pair is further disrupted by steric interference with a conserved palm domain loop C-terminal to a helix of the fingers domain, which acts as a separation wedge (Figure 1d). Engagement of DNA in the active site therefore results in melting of two downstream base pairs.

Importantly, the upstream duplex geometry and thumb positioning are preserved relative to the primer–template complex (Figure 1c), indicating that strand displacement does not require a distinct global polymerase conformation. The palm domain loop that acts as the separation wedge contacts the downstream duplex in the same configuration observed during gap-filling synthesis, is conserved across Pol δ homologues, and is structurally conserved in Pol α, which lacks strand displacement activity^18^ (Figure 1f). These observations demonstrate that the separation wedge is a constitutive feature of the B-family fold rather than a structural element dedicated to or induced by strand displacement. The energetic cost of each strand displacement cycle is higher than gap-filling synthesis, as forward translocation must be coupled to melting of downstream base pairs rather than reading a pre-exposed single-stranded template. This intrinsic energetic penalty explains why strand displacement activity in human Pol δ is weak and clamp-dependent^19,20^: PCNA engagement provides the mechanical continuity required for Pol δ to overcome the barrier of downstream base-pair melting.

### Toolbelt assembly pre-positions FEN1 for substrate transfer

To capture the strand-displacement machinery in a clamp-mediated toolbelt configuration, we assembled Pol δ–PCNA with FEN1 on the flap DNA substrate using a chain-terminated primer in the presence of dTTP. Cryo-EM reconstruction yielded a 3.5 Å map resolving all components of the complex (Figure 2a-b; Extended Data Figure 2; Extended Data Table 2). The architecture is consistent with a toolbelt arrangement in which Pol δ and FEN1 occupy distinct PIP-binding sites on PCNA (Figure 2a).

**Figure 2.**
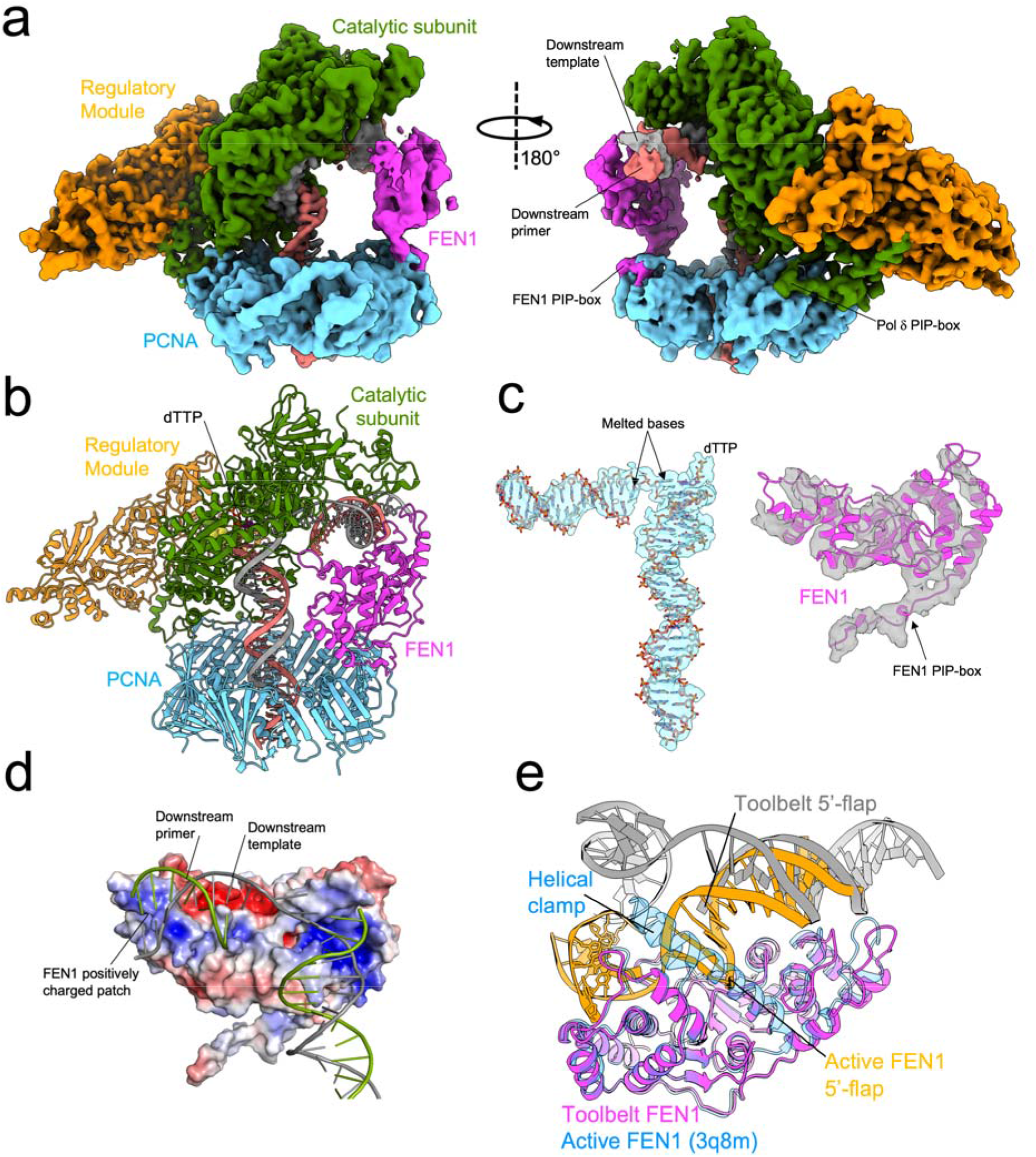
Structure of the Pol δ–flap DNA–dTTP–PCNA–FEN1 toolbelt. **(a)** Two views of the cryo-EM map of the toolbelt, with components colour-coded. **(b)** Atomic model of the toolbelt, positioning FEN1 adjacent to the elongating polymerase. **(c)** Cryo-EM density and fitted model of the flap DNA (left) and FEN1 (right) from the reconstruction shown in (a). **(d)** Electrostatic surface representation of FEN1 highlighting a positively charged patch that engages the downstream duplex of the flap substrate. **(e)** Superposition of FEN1–DNA from the toolbelt complex with the X-ray structure of FEN1 bound to flap DNA^21^ (PDB: 3Q8M), showing that the positions of the 5′ flap are aligned, suggesting that FEN1 in the toolbelt is prepositioned to accept and process the substrate.

The primer terminus remains engaged in the polymerase active site with bound dTTP, closely matching the strand displacement structure lacking FEN1 (Figures 2b-c and Figure 1). The upstream duplex and thumb positioning are preserved, indicating that FEN1 recruitment does not perturb the engaged polymerase conformation. The downstream duplex remains loosely tethered to the Pol δ N-terminal domain and extends toward FEN1 (Figure 2b), with two base pairs melted at the flap junction (Figure 2c). Although density at the FEN1–DNA interface does not permit resolution of atomic contacts, the duplex consistently localises against a positively charged surface patch of FEN1 implicated in duplex engagement (Figure 2d), with reduced local map resolution over FEN1 indicating partial flexibility within the complex (Figure 2c; Extended Data Figure 2). In contrast to the complex without FEN1, all three PCNA monomers are well defined (Figure 2a), suggesting that FEN1 engagement with the clamp — and its partial contact with the downstream duplex — contributes to ring ordering. Superposition of FEN1 from the toolbelt complex with the crystal structure of FEN1 bound to flap DNA (PDB: 3Q8M)^21^ shows that the 5′ flap ends are aligned (Figure 2e), indicating that FEN1 in the toolbelt is pre-positioned to accept and process the substrate without requiring large-scale repositioning upon polymerase disengagement.

A previous cryo-EM structure of the human Pol δ–PCNA–FEN1 toolbelt was assembled on primer–template DNA lacking a downstream flap ^16^, and therefore could not address the spatial relationship between the FEN1 active site and the displaced strand. The present structure directly visualises the trajectory of the downstream duplex toward the FEN1 active site cleft in the context of an actively synthesizing, strand-displacement-competent holoenzyme, establishing that the toolbelt is a geometrically pre-organised assembly rather than a passive co-occupancy arrangement.

### An exchange-prone configuration of Pol ***δ*** licenses DNA handoff to FEN1

The dTTP-trapped toolbelt structure captures Pol δ in the engaged, catalytically competent state, with the primer terminus stably anchored in the active site. In this configuration, the DNA substrate is not accessible to FEN1, and substrate handoff cannot occur. We therefore asked what drives the transition from this engaged state to one permissive for FEN1 capture. We reasoned that in the absence of incoming nucleotide — as occurs transiently between incorporation events during strand displacement synthesis — the polymerase may adopt an open, exchange-prone configuration with reduced grip on the primer terminus, generating a geometry permissive for FEN1 access.

To test this, we assembled Pol δ and PCNA on the flap-containing substrate in the absence of incoming nucleotide, added FEN1, separated the complex by gel filtration, and analysed the complex fraction by cryo-EM (Figure 3a; Extended Data Figure 3). In contrast to the homogeneous toolbelt population observed in the presence of dTTP (Figure 3b), reference-free 2D classification revealed multiple particle populations corresponding to a Pol δ–PCNA–DNA–FEN1 toolbelt assembly, dissociated Pol δ, and intact FEN1–PCNA–DNA complexes (Figure 3a). Toolbelt particles yielded a 3.4 Å reconstruction in which Pol δ and FEN1 simultaneously bind PCNA, while the upstream duplex remains tethered to the polymerase yet disengaged from the active site (Figure 3c-d; Extended Data Figure 3; Extended Data Table 3). In this assembly, Pol δ adopts an exchange-prone configuration characterised by open fingers and a retracted thumb (Figure 3d). The downstream duplex trajectory was modelled by flexible fitting with distance and base-pair restraints guided by the weak cryo-EM density, consistent with Pol δ bending the flap junction prior to engaging the primer-template in the active site (Figure 3c). Notably, FEN1 exhibits reduced local resolution and is visible only in the unsharpened map (Figure 3c, Extedended Data Figure 3), pointing to increased flexibility in this exchange-prone configuration.

**Figure 3.**
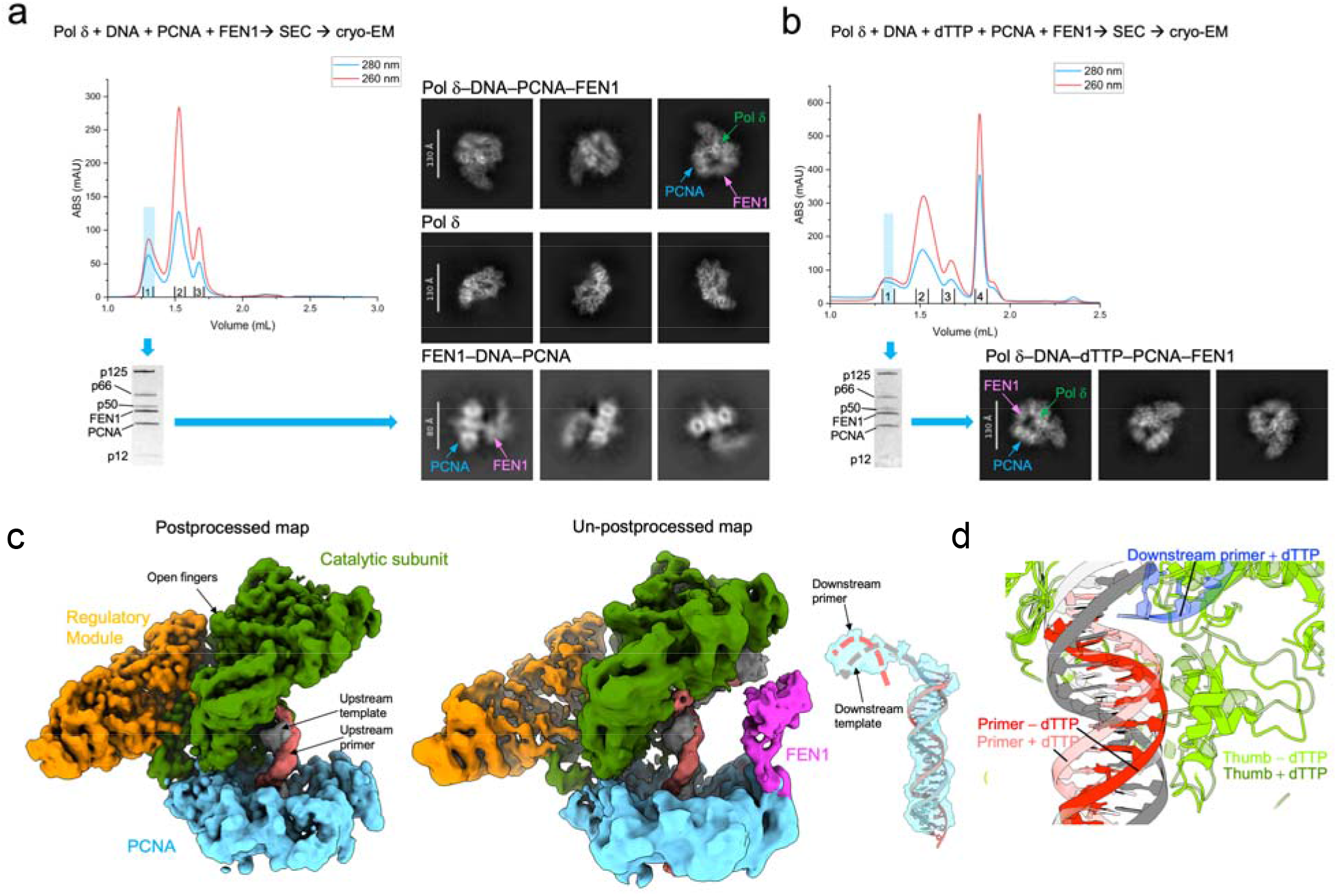
Structural basis of substrate transfer from Pol δ to FEN1. **(a)** Size-exclusion chromatography purification of the Pol δ–flap DNA–PCNA–FEN1 complex in the absence of dTTP. SDS–PAGE of the peak fraction shows all components of the complex, and this fraction was used for cryo-EM analysis. Reference-free 2D class averages reveal a mixed population of particles corresponding to the intact complex, free Pol δ, and FEN1–DNA–PCNA complexes, indicating that substrate transfer to FEN1 is accompanied by dissociation of Pol δ from PCNA. **(b)** Size-exclusion chromatography purification of the Pol δ–flap DNA–PCNA–FEN1 complex in the presence of dTTP and cryo-EM analysis, carried out under the same conditions as in (a). Reference-free 2D class averages reveal a homogeneous population of particles corresponding to the intact toolbelt complex, indicating that nucleotide addition stabilises the complex and prevents partitioning. **(c)** Postprocessed and un-postprocessed cryo-EM maps of the Pol δ–flap DNA–PCNA–FEN1 complex without dTTP. DNA remains bound to Pol δ but is disengaged from the active site. FEN1 density is visible only in the un-postprocessed map, consistent with increased flexibility of this component. The upstream duplex is well resolved, whereas the downstream duplex is flexible and poorly defined. The position of the downstream duplex shown as a dashed model was obtained by restrained flexible fitting with base-pair and distance restraints to residual density, and is presented to illustrate the probable trajectory of the substrate. **(d)** Structural overlay of the toolbelt complexes with and without dTTP, aligned on the catalytic domain, highlighting divergent positions of the DNA and thumb domain.

These observations indicate that the exchange-prone configuration of Pol δ generates a geometry permissive for FEN1 access while the duplex remains topologically confined within PCNA. The coexistence of toolbelt particles together with dissociated Pol δ and intact FEN1–PCNA–DNA assemblies in the same dataset suggests that the clamp-mediated complex formed during purification subsequently partitions into multiple assemblies. In this model, release of the primer terminus from the polymerase active site allows FEN1 to capture the emerging flap while both enzymes remain tethered to PCNA. Geometric analysis of the toolbelt structure and post-handoff FEN1–PCNA–DNA structure (presented below) shows that the upstream duplex, already threaded through the PCNA ring, requires only a ∼30° rotation to reach the FEN1-engaged position, rendering intramolecular handoff strongly favoured over an intermolecular mechanism that would require the DNA to exit the ring and reassemble. Consistent with this interpretation, FEN1–PCNA–DNA particles are observed exclusively in the nucleotide-free dataset — not when dTTP is present — despite free components being equally available under both conditions, demonstrating that partitioning is gated by the catalytic state of Pol δ rather than by component availability. Following substrate transfer, Pol δ is no longer anchored to the primer terminus and its interaction with the clamp becomes labile, consistent with the population of dissociated polymerase particles observed in the 2D class averages (Figure 3a).

### Structure of FEN1–PCNA engaging flap DNA

A 3.9 Å cryo-EM reconstruction of FEN1 bound to flap DNA and PCNA resolves a state consistent with a post-polymerase handoff (Figure 4a; Extended Data Figure 4; Extended Data Table 4). FEN1 engages one monomer of the PCNA ring via its PIP box, leaving the remaining two PIP-binding sites unoccupied (Figure 4a), consistent with previous structural studies^10,22–24^.

**Figure 4.**
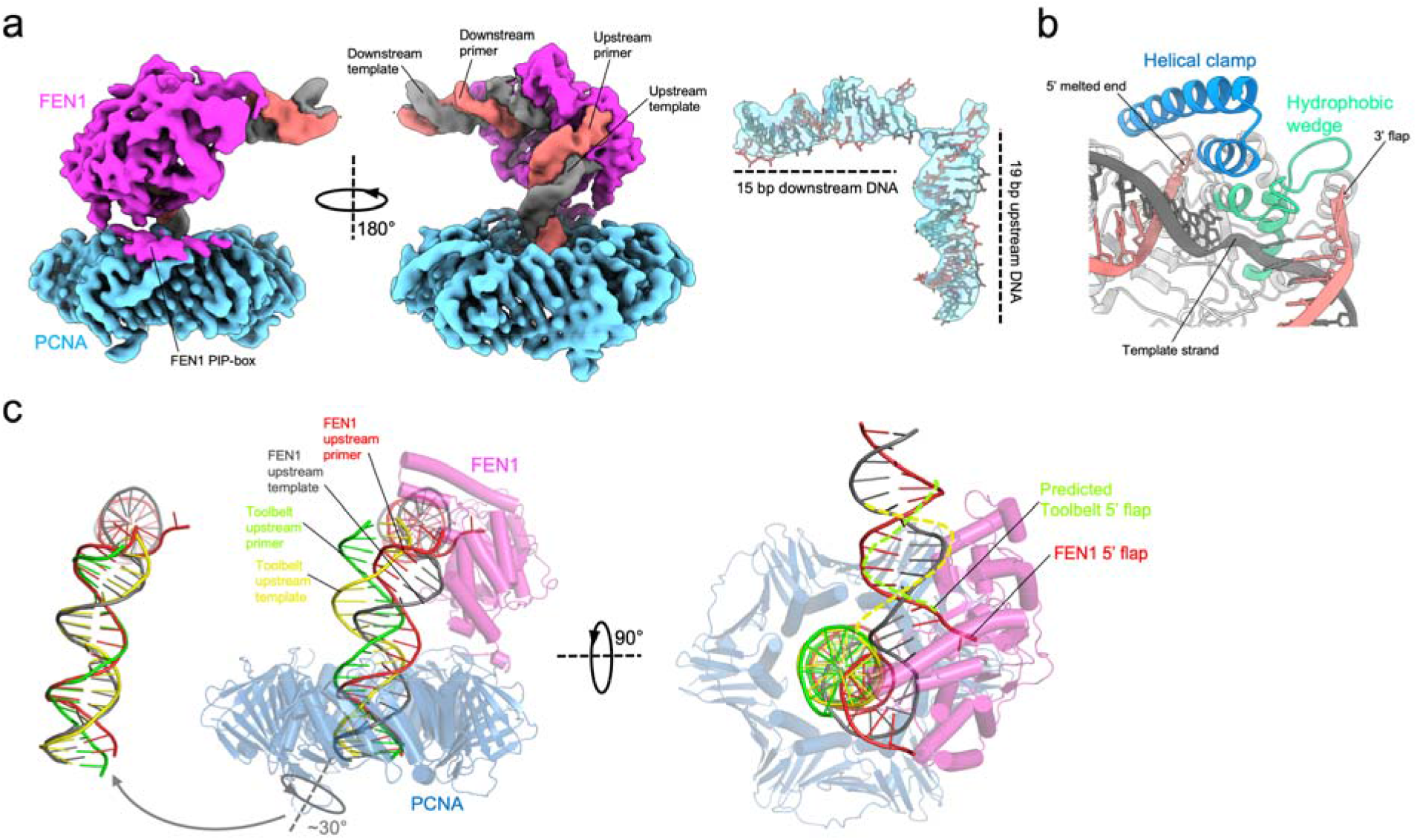
Structure of FEN1 bound to PCNA and flap DNA. **(a)** Two views of the cryo-EM reconstruction. The inset shows the map and model of the DNA substrate, highlighting the L-shaped DNA conformation induced by FEN1 engagement. **(b)** Close-up of the FEN1–DNA–PCNA model showing the flap junction positioned in the FEN1 active site. The terminal base pair at the 5′ end is melted and the flap overhang is not resolved, consistent with the post-cleavage state reported previously^21^ (PDB: 3Q8K. **(c)** Two views of the overlay, aligned on PCNA, of the FEN1–DNA–PCNA structure with the Pol δ–flap DNA–PCNA–FEN1 toolbelt model without dTTP. For clarity, only the DNA from the toolbelt structure is shown. The inset to the left shows that the upstream duplex in the two structures differs by a simple ∼30° rotation about an axis threading the PCNA ring. Although the downstream duplex cannot be unambiguously modelled, the approximate position of its 5′ flap inferred from the fitted model aligns closely with the 5′ flap observed in the FEN1 structure, suggesting that FEN1 captures the 5′ flap directly from the toolbelt configuration without clamp dissociation.

Upon binding FEN1, the DNA adopts an elbow-shaped conformation (Figure 4a). The template strand remains continuous, while the complementary strands are interrupted within the nuclease active site, producing an approximately 90° bend. Both terminal regions of the substrate adopt double-stranded conformations: the downstream duplex engages FEN1 exclusively, whereas the upstream duplex engages FEN1 and passes through the PCNA ring (Figure 4a).

The 5’-end of the downstream complementary strand is positioned at the FEN1 catalytic center (Figure 4b). The ssDNA flap is unseen and the terminal 5’ nucleotide is flipped and melted (Figure 4a-b), recapitulating the post-cleavage configuration observed in a prior crystal structure^21^. Despite the use of the catalytically impaired D181A mutant, this indicates that cleavage occurred during sample preparation, consistent with its residual catalytic activity ^25^. At the 3’ end of the upstream complementary strand, a single overhanging nucleotide is visible, stabilized by the hydrophobic wedge of FEN1 (Figure 4b). Multi-body refinement reveals substantial rotational flexibility of the FEN1–DNA module relative to PCNA (Extended Data Figure 4), consistent with prior observations of clamp-associated nuclease mobility in native complexes^24^.

We next compared the FEN1-engaged structure with the exchange-prone toolbelt assembly. Flexible fitting of the toolbelt downstream duplex defines a trajectory for the emerging 5’ flap that aligns with the path observed in the FEN1-engaged structure (Figure 4c). The upstream duplexes are approximately related by a ∼30° rotation about an axis encompassing the PCNA ring (Figure 4c), whereas the downstream duplex requires additional rearrangements to reach the FEN1-engaged configuration. While intermediate states are not directly resolved, these observations are consistent with a model in which the transition to the FEN1-engaged state involves rotation of the clamp-bound DNA without release of PCNA, linking substrate transfer to the exchange-prone Pol δ configuration.

## Discussion

### Structural basis of strand displacement across polymerase families

Strand displacement by human Pol δ emerges from structural elements intrinsic to the B-family polymerase fold rather than from a dedicated separation machinery. The downstream duplex follows the trajectory defined by palm domain contacts that position unpaired template bases during processive synthesis, with the complementary strand arranged as a geometric consequence. At the duplex junction, a conserved palm loop acts as a passive wedge that destabilises the first downstream base pairs, while R224/R225 in the N-terminal domain extend the templating path into the duplex (Figure 5a). The presence of this loop in Pol α, which lacks strand displacement activity^18^, indicates that it is not sufficient to confer this function. Instead, PCNA engagement^19,20^ enables Pol δ to overcome the energetic barrier of downstream base-pair melting within an otherwise constitutive structural framework.

**Figure 5.**
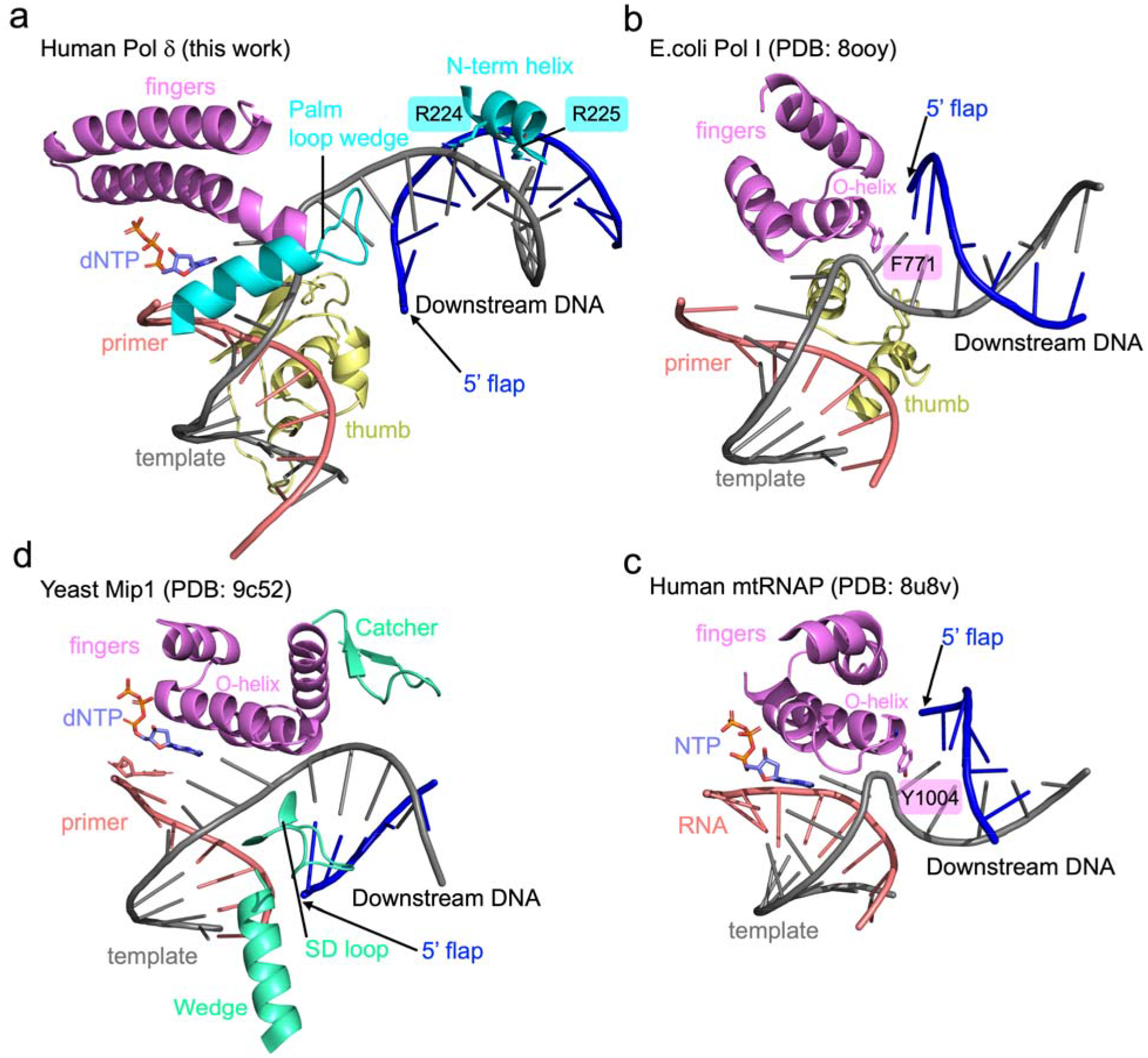
Structural diversity of strand displacement mechanisms across polymerase families. **(a)** Structure of human Pol δ engaged on flap DNA from this work. The palm loop wedge (cyan) destabilises the first downstream base pair, while R224 and R225 on the N-terminal helix extend the templating path into the downstream duplex, directing the 5′ flap away from the thumb domain. **(b)** Structure of *E. coli* Pol I engaged on flap DNA^12^ (PDB: 8ooy). The O-helix of the fingers domain positions F771 as a separation pin that wedges apart downstream base pairs, directing the 5′ flap toward the thumb domain. **(c)** Structure of human mitochondrial RNA polymerase^26^ (mtRNAP) engaged on a DNA–RNA hybrid (PDB: 8u8v). Y1004 on the O-helix acts as a separation pin analogous to Pol I F771. **(d)** Structure of yeast mitochondrial polymerase Mip1 engaged on flap DNA^13^ (PDB: 9c52). Despite belonging to the A-family, Mip1 directs the 5′ flap away from the thumb, as in Pol δ, achieved through a lineage-specific SD loop and wedge insertion rather than a separation pin, reflecting partial convergence on the B-family architecture. In all panels, the template strand is grey, the primer/RNA strand is salmon, and the downstream primer is in dark blue.

Strand displacement thus represents a functional outcome realised through distinct structural solutions across polymerase families. In A-family *E.coli* Pol I^12^, similar to human mitochondrial RNAP^26^, a conserved separation pin in the fingers domain wedges apart downstream base pairs along a trajectory directed toward the thumb (Figure 5b-c). In contrast, the yeast mitochondrial polymerase Mip1^13^, although an A-family enzyme, directs the downstream 5′ end away from the thumb, as in Pol δ, and achieves strand separation through conserved elements augmented by a lineage-specific insertion, reflecting partial convergence on the B-family architecture (Figure 5d).

The divergence between Pol I and Pol δ has direct implications for Okazaki fragment maturation. When aligned on the upstream duplex, the 5′ flap exits in opposite directions in the two systems (Figure 5a-b). In Pol I, the nuclease is fused in cis and accesses the flap via a flexible linker^12^. In Pol δ, FEN1 is recruited in trans to the front face of PCNA, precisely where the flap emerges, indicating that clamp-mediated toolbelt assembly is dictated by the structural logic of the B-family fold.

### Toolbelt framework for eukaryotic Okazaki fragment maturation

Eukaryotic Okazaki fragment maturation requires the coordinated action of Pol δ, FEN1 and Lig1, all recruited to the PCNA ring. The concept that a sliding clamp can simultaneously accommodate multiple client enzymes on distinct subunits was first established in the archaeon *Sulfolobus solfataricus*, whose heterotrimeric PCNA presents a dedicated binding site for each of the three OFM enzymes^27^. The structural basis for subunit-specific client recognition was subsequently revealed by crystal structures of binary PCNA–enzyme subcomplexes^28^, and reconstitution experiments confirmed that the heterotrimer coordinates sequential DNA synthesis, flap removal and ligation in a single assembly^29^. Whether an analogous toolbelt operates in eukaryotes, where PCNA is a homotrimer with equivalent subunits, has been debated, with sequential switching and toolbelt models both proposed^8,9,15^. The structures reported here resolve this question and define a mechanistic framework for the eukaryotic maturation cycle.

When Pol δ−PCNA arrives at the nick of a downstream Okazaki fragment, the polymerase must incorporate at least one nucleotide to generate a 5’ flap before FEN1 can engage^21^. Our strand displacement structure captures this minimal state: a single dTTP incorporation has melted two downstream base pairs, generating a 2-nucleotide 5’ flap sufficient for FEN1 recognition. The dNTP-trapped toolbelt structure shows that FEN1 can be recruited to PCNA while Pol δ remains engaged in strand displacement synthesis, demonstrating that simultaneous occupancy is compatible with an active polymerase. In this state, the 5’ end of the downstream duplex is oriented toward the FEN1 active site but the substrate remains anchored in Pol δ, precluding transfer. The exchange-prone toolbelt without dNTP captures the subsequent transient state in which release of the primer terminus weakens Pol δ engagement, poising the complex to partition into dissociated Pol δ and a FEN1–PCNA–DNA intermediate. A simple rotation of the upstream duplex defines the transfer trajectory, pivoting the DNA about a PCNA-encompassing axis to reposition the flap from Pol δ to FEN1 without clamp dissociation. This motion is enabled by the topological confinement of DNA within PCNA.

These observations support a toolbelt model in which Pol δ and FEN1 are simultaneously bound to PCNA during substrate transfer, with exchange triggered by polymerase disengagement rather than competitive displacement. Earlier biochemical experiments in the yeast system using engineered PCNA heterotrimers demonstrated that sequential partner switching on a single PCNA monomer is sufficient to support complete Okazaki fragment maturation, albeit at lower rates^15^. However, later kinetic studies showed that the PCNA–Pol δ–FEN1 complex moves processively through iterative nick translation cycles^9^, with FEN1 remaining stably associated with PCNA between successive cleavage events, consistent with our data.

Our data supporting Pol δ dissociation from PCNA upon substrate transfer, and stable engagement of cleaved DNA within FEN1, provides a structural explanation for previous biochemical observations that nick translation is kinetically inefficient in the human system^8^: Pol δ poorly re-engages substrates once captured and cleaved by FEN1, whereas Lig1 promotes product release to drive ligation^8^. Our previous structure of Lig1 engaging nicked DNA and bound to PCNA together with FEN1 captured the post-handoff ligation toolbelt^10^, completing the maturation cycle. Consistent with this, direct affinity measurements showed that Pol δ binding to PCNA-loaded DNA weakens ∼10-fold as the substrate transitions from a 2-nt gap to a 5′ flap^8^, and both the FEN1−Pol δ−PCNA and FEN1−Lig1−PCNA toolbelt assemblies were directly detected biochemically^8^, corroborating our structural data. We note that recent studies from our laboratories ^10,16^ and others^24,30,31^ report that toolbelt arrangements with two enzymes bound to a single clamp ring are increasingly observed across organisms, suggesting that simultaneous clamp occupancy by consecutive enzymes is a conserved organisational principle of DNA metabolism.

Taken together, our data show that PCNA serves as a topological organiser that maintains the DNA substrate throughout the Okazaki fragment maturation cycle, a geometric scaffold that pre-positions each incoming enzyme adjacent to the active site of its predecessor, and a conformational gating platform that couples substrate transfer to the catalytic state of the polymerase. In this framework, the clamp ring is not merely a processivity factor but the structural logic that makes coordinated, directional maturation possible.

## Materials and Methods

### Protein expression and purification

Full-length human PCNA (UniProt: P12004) was cloned in pETDuet-1 MCS1 (Novagen) Amp+ to obtain 6x His N-terminally-tagged protein. Wild-type PCNA was expressed and purified as described previously^16^. Briefly, protein expression was carried out in 2YT medium supplemented with ampicillin. Cultures were grown at 37 degrees C to an OD600 of approximately 1.2, induced with 0.5 mM IPTG, and further incubated at 16 degrees C for 19 h. Cells were harvested by centrifugation, resuspended in lysis buffer, and lysed by lysozyme treatment followed by sonication. All subsequent purification steps were performed at 4 degrees C. Insoluble material was removed by high-speed centrifugation, and the clarified lysate was loaded onto a HisTrap HP affinity column (Cytiva) pre-equilibrated in binding buffer. After extensive washing, bound proteins were eluted using an imidazole gradient and further purified by anion-exchange chromatography on a HiTrap Q HP column (Cytiva). Final purification was achieved by size-exclusion chromatography on a Superdex 200 column (Cytiva) equilibrated in storage buffer. Fractions containing pure PCNA were pooled, concentrated, flash-frozen in liquid nitrogen and stored at -80 degrees C.

Human Pol δ wild-type and D515V mutant were expressed as described previously^8,16^. Briefly, a MultiBac expression plasmid encoding all four subunits of Pol δ was introduced into DH10MultiBac cells to generate bacmid DNA. This bacmid was then transfected into Sf9 insect cells using FuGENE HD, following the manufacturer’s protocol, to produce recombinant baculovirus. The virus was amplified twice to obtain a high-titer P3 viral stock. Pol δ was expressed by infecting a 4 L Sf9 suspension culture at a cell density of 2 x 10^6 cells/ml with the P3 virus and incubated for 70 hours. Cells were harvested and the clarified lysate was first purified using a HisTrap HP affinity column (Cytiva) under low-salt conditions, followed by anion-exchange chromatography on a Mono Q column (Cytiva). Fractions containing all four Pol δ subunits were further purified by size-exclusion chromatography on a Superdex 200 column (Cytiva). The purified protein was concentrated, pooled, flash-frozen, and stored at -80 degrees C.

D181A FEN1 was expressed and purified as described previously^32^. Briefly, the plasmid was transformed into E. coli BL21(DE3) cells and cultured in 2YT media. Expression was induced by the addition of 0.2 mM IPTG when the OD600 of the culture reached 0.8. Cells were harvested and lysed in buffer A [50 mM Tris-HCl pH 7.5, 5% (v/v) glycerol, 750 mM NaCl, 10 mM beta-mercaptoethanol (BME), and 30 mM imidazole] by a combination of lysozyme treatment and sonication. Purification was carried out by two sequential Ni-NTA columns, separated by SUMO protease cleavage. Proteins were then concentrated and further purified over HiLoad Superdex-75 pg size exclusion columns using a buffer containing 50 mM HEPES-KOH pH 7.5, 500 mM NaCl, 2 mM dithiothreitol (DTT), and 10% (v/v) glycerol. Fractions containing pure D181A FEN1 were collected, flash frozen, and stored at -80 degrees C.

### DNA substrates

All oligonucleotides were obtained from Integrated DNA Technologies (IDT) and supplied with HPLC purification, except where noted. The flap DNA substrate used for the strand-displacement Pol δ holoenzyme and Pol δ–PCNA–FEN1 toolbelt complexes was assembled from three oligonucleotide strands: a template strand (5′-GGTGATCGTTCGCTACAAGTCGTCAGGATTCCAGGCAGGCTAT-3′), a primer strand (5′-ATAGCCTGCCTGGAATCCTGACGAddC-3′, where ddC denotes a 3′ dideoxy chain terminator), and a flap strand (5′-AGACACATGGTTGTAGCGAACGATCACC-3′). Annealing reactions were performed in a buffer containing 50 mM Tris-HCl (pH 7.5) and 50 mM NaCl. Equimolar concentrations of the three oligonucleotides were combined, heated to 95 °C, and gradually cooled to room temperature at a rate of 2 °C min ¹ using a thermocycler.

The flap DNA substrate used for the FEN1–DNA–PCNA complex was assembled from three oligonucleotide strands: a flap strand (5′-AGACACATGGATGTAGCGAACGATCACC-3′), a template strand (5′-GGTGATCGTTCGCTACATGTCGTCAGGATTCCAGGCAG-3′), and a complementary strand (5′-CTGCCTGGAATCCTGACGACT-3′). Oligonucleotides were obtained from Eurofins. Annealing reactions were performed in a buffer containing 20 mM Tris-HCl (pH 7.5) and 25 mM NaCl. Equimolar concentrations of the three strands were combined, heated to 92 °C for 2 minutes, and allowed to cool gradually to room temperature.

### Cryo-EM sample preparation

For the strand displacement complex, Pol δ and PCNA were mixed at a 1:1 molar ratio (Pol δ:PCNA trimer) with the three-strand flap substrate at a 1:1 molar ratio (Pol δ:DNA) in assembly buffer (25 mM HEPES pH 7.5, 100 mM KAc, 10 mM CaCl2, 0.02% (v/v) NP-40, 1 mM DTT). The complex was purified by gel filtration on a Superose 6 Increase 3.2/300 column (Cytiva) equilibrated in assembly buffer without nucleotide. Peak fractions corresponding to the Pol δ–PCNA–DNA complex were collected. Immediately before vitrification, dTTP was added to a final concentration of 250 μM and the sample was incubated for 7 min on ice. Cryo-EM grids (Quantifoil R2/1 Cu 300 mesh with a 2 nm continuous carbon support layer) were glow-discharged for 10 s at 10 mA and 3 μl of sample was applied. Grids were blotted for 2 s at 4 °C and 100% humidity and plunge-frozen in liquid ethane using a Vitrobot Mark IV (Thermo Fisher Scientific).

For the dTTP-trapped toolbelt complex (Pol δ–PCNA–FEN1 with dTTP), Pol δ, PCNA, and FEN1 D181A were mixed at a 1:1:2 molar ratio with the three-strand flap substrate (chain-terminated primer) in assembly buffer containing 200 μM dTTP. The complex was purified by gel filtration as above. Peak fractions were collected and vitrified as described above without additional nucleotide addition.

For the nucleotide-free toolbelt complex (exchange-competent state), Pol δ and PCNA were first assembled on the flap substrate as described above. FEN1 D181A was then added at a 2-fold molar excess over Pol δ and the mixture was purified by gel filtration n. Peak fractions were collected and vitrified immediately.

The FEN1–DNA–PCNA complex was separated by size-exclusion chromatography using a Superdex 200 Increase 3.2/300 microSEC column (GE Life Sciences). A 50 μl aliquot of the following mixture—4 μM FEN1 (D181A), 5 μM DNA, and 5 μM PCNA trimer in 0.1 mM ATP, 25 mM HEPES (pH 7.5), 100 mM KAc, 10 mM MgCl, and 0.5 mM TCEP—was loaded onto the column pre-equilibrated in the same buffer. UltrAuFoil R1.2/1.3 300-mesh gold grids were glow discharged at 40 mA for 5 min using a Quorum GloQube and coated with a layer of graphene oxide (Sigma). A 3 μl aliquot of the selected fraction from the size-exclusion chromatography was applied to the grid. The grid was then plunge-frozen into liquid ethane using a Vitrobot Mark IV (FEI/Thermo Fisher Scientific) set to 4 °C and 100% relative humidity, with a blot force of 10, a wait time of 0 s, and a blot time of 3 s.

### Cryo-EM data collection

Cryo-EM datasets for the strand-displacement Pol δ complex (12,194 movies), toolbelt complex with dTTP (16,393 movies), and toolbelt complex without dTTP (12,582 movies) were collected on a Titan Krios G4 (Thermo Fisher Scientific) operating at 300 kV, equipped with a Falcon 4i direct electron detector in counting mode. All datasets were acquired at a nominal magnification of 165,000×, corresponding to a calibrated pixel size of 0.76 Å/pixel, with a defocus range of −3.0 to −1.0 μm in 0.4 μm steps. Data collection was performed in EPU using aberration-free image shift (AFIS). Specimen dose rates were 5.96, 6.05, and 6.26 e /pixel/s for the strand-displacement, toolbelt with dTTP, and toolbelt without dTTP datasets, respectively. FEN1–DNA–PCNA dataset: data were collected on a Titan Krios G3 (Thermo Fisher Scientific) operating at 300 kV at the Midlands Regional Cryo-EM facility (Leicester Institute for Structural and Chemical Biology). Movies (2,150) were recorded in super-resolution mode using a Gatan K3 direct electron detector at a dose rate of approximately 16.5 e /pixel/s.

### Cryo-EM data processing

Strand-displacement dataset: A total of 12,194 raw movies were imported into RELION 5.0b and subjected to beam-induced motion correction using the RELION implementation of MotionCor2^33^, with all frames retained and patch alignment set to 4 × 4. Motion-corrected micrographs were imported into cryoSPARC^34^ for CTF estimation using Patch CTF. Particles were initially picked using a blob-based approach, yielding 1,874,538 candidates extracted with a box size of 400 pixels. Three rounds of 2D classification reduced this set to 165,407 particles, which were used to train a Topaz neural network picker. The trained topaz model was used to re-pick and re-extract the full dataset, and four subsequent rounds of 2D classification produced a curated set of 262,654 particles. These were exported to RELION^35^ for two rounds of 3D classification, generating four distinct classes. The 139,707 particles comprising the best-resolved class were subjected to 3D refinement with spIsoNet misalignment correction^36^, yielding an initial reconstruction at 3.5 Å. Application of spIsoNet anisotropy correction improved the resolution to 3.1 Å. To assess conformational heterogeneity, 3D variability analysis (3DVA)^17^ was performed on the final particle set. The consensus map was sharpened and post-processed with EMReady^37^. Toolbelt with dTTP dataset: A total of 16,393 EER-format movies were motion-corrected using Warp (v1.0.9)^38^, with movies fractionated into 40 frames at a per-frame dose of 1 e /Å², processed at the original pixel size with a B-factor of −150 Å² for dose weighting. Motion-corrected micrographs were imported into cryoSPARC^34^ for CTF estimation using CTFFIND4. Particles were initially picked using a blob-based approach, yielding 2,558,694 particles extracted with a box size of 400 pixels. Two rounds of 2D classification retained 12,646 particles, which were used to train a Topaz neural network picker. The trained model was used to re-pick and re-extract the dataset, and five subsequent rounds of 2D classification yielded 57,254 particles, which served as templates for a subsequent round of template-based picking. After re-extraction, one round of 2D classification, four rounds of heterogeneous refinement, and three rounds of 2D classification produced a curated set of 200,963 particles. These were exported to RELION^35^ for two rounds of 3D classification, generating four distinct classes. The 148,674 particles from the best-resolved class were imported into cryoSPARC for non-uniform refinement, yielding an initial reconstruction at 3.6 Å. Application of spIsoNet anisotropy correction improved the resolution to 3.5 Å. The consensus map was sharpened and post-processed with EMReady^37^. Toolbelt without dTTP dataset: A total of 12,582 EER-format movies were motion-corrected using Warp (v1.0.9)^38^ with the same fractionation and dose-weighting parameters as the toolbelt dataset. Motion-corrected micrographs were imported into cryoSPARC^34^ for CTF estimation using CTFFIND4. Particles were initially picked using the Topaz model trained on the toolbelt dataset, yielding 2,474,116 candidates extracted with a box size of 400 pixels. One round of 2D classification, three rounds of heterogeneous refinement, and two additional rounds of 2D classification produced a curated set of 129,184 particles. These were exported to RELION for one round of 3D classification, generating four distinct classes. The 105,913 particles from the best-resolved class were imported into cryoSPARC for non-uniform refinement, yielding a reconstruction at 3.4 Å that was unchanged after spIsoNet anisotropy correction. The consensus map was sharpened and post-processed with EMReady^37^. FEN1–DNA–PCNA dataset: Image processing was carried out in RELION 3.1^35^. Movie stacks were imported and corrected for beam-induced motion using MotionCor2^33^. Contrast transfer function (CTF) parameters for each micrograph were estimated using CTFFIND4. Particle picking was performed using the external crYOLO^39^ picker. Particles were extracted with a box size of 258 pixels. Subsequent processing included 2D classification, 3D classification, and 3D refinement, followed by rounds of Bayesian polishing, CTF refinement, and multibody refinement. Resolution of the final reconstruction at 3.8 Å. The final map was sharpened and post-processed using EMReady^37^.

### Model building

Model building of the strand-displacement Pol δ holoenzyme: the initial model was the ternary complex (PDB ID: 6TNY), docked into the cryo-EM density. The downstream DNA duplex was built de novo in Chimera^40^ and the complete model refined through iterative flexible fitting in ISOLDE^41^ followed by real-space refinement in Phenix^42^ using geometric restraints. Model building of the Pol δ–PCNA–FEN1 complex bound to flap DNA and dTTP: the initial model was the Pol δ–PCNA–FEN1 ternary assembly bound to P/T DNA (PDB ID: 6TNZ). The downstream duplex DNA was built in Chimera and refined as above using ISOLDE and Phenix with geometric restraints. For the Pol δ–PCNA–FEN1 complex without dTTP, model building followed the same procedure; however, due to weak density for the downstream duplex, only the upstream DNA segment was modelled and deposited. Model building of the FEN1– PCNA–flap DNA complex: initial models comprised PCNA from 6TNY 18 and FEN1–DNA from PDB 3Q8K 14. B-form DNA was extended to model the upstream duplex, and the complete assembly refined using ISOLDE and Phenix with geometric restraints.

## Data availability

Cryo-EM maps and atomic coordinates have been deposited in the Electron Microscopy Data Bank (EMDB) and the Protein Data Bank (PDB) under accession codes to be assigned upon acceptance. All other data supporting the findings of this study are available from the corresponding authors upon reasonable request.

Maps for the flap DNA, PCNA, Pol δ in strand displacement, FEN1 toolbelt-engaged, and FEN1 toolbelt-disengaged complexes, along with the FEN1–PCNA map have been deposited in the EMDB with the accession codes EMD-80308, EMD-80285, EMD-80304, EMD-80307, respectively. The corresponding atomic models were deposited in the Protein Data Bank with the accession codes 25QR, 25PJ, 25QN, 25QQ. The 3DVA maps of frames 1 and 20 of the first component from strand-displaced DNA were deposited as EMD-80716 and EMD-80714, respectively. The corresponding atomic models were deposited in the Protein Data Bank with the accession codes 26KK and 26KG, respectively.

## Author Contributions

I.Z. and A.U.D. carried out cryo-EM sample preparation, data collection, image processing, and atomic model building of Pol δ and FEN1 complexes. M.T. expressed and purified the recombinant proteins. L.Z. assisted with cryo-EM data collection. K.B. solved the structure of the FEN1−DNA−PCNA complex. R.A. helped with cryo-EM data analysis and V-S. R. helped with the overall analysis. S.M.H. and A.D.B. supervised research. A.D.B wrote the paper with the help of the other authors.

## Acknowledgements

This research was supported by King Abdullah University of Science and Technology (KAUST) through core funding (to S.M.H. and A.D.B.) and by the Competitive Research Award Grant CRG8 URF/1/4036-01-01. We thank the KAUST Imaging and Characterization Core Labs for access to cryo-EM infrastructure and support with data collection. We thank members of the Hamdan and De Biasio laboratories for helpful discussions.

