## Supplementary Information File for "Structural basis of nick translation in human DNA replication"

**SI Guide**

**Contents**

- Extended Data Figures 1–4
- Extended Data Tables 1–4

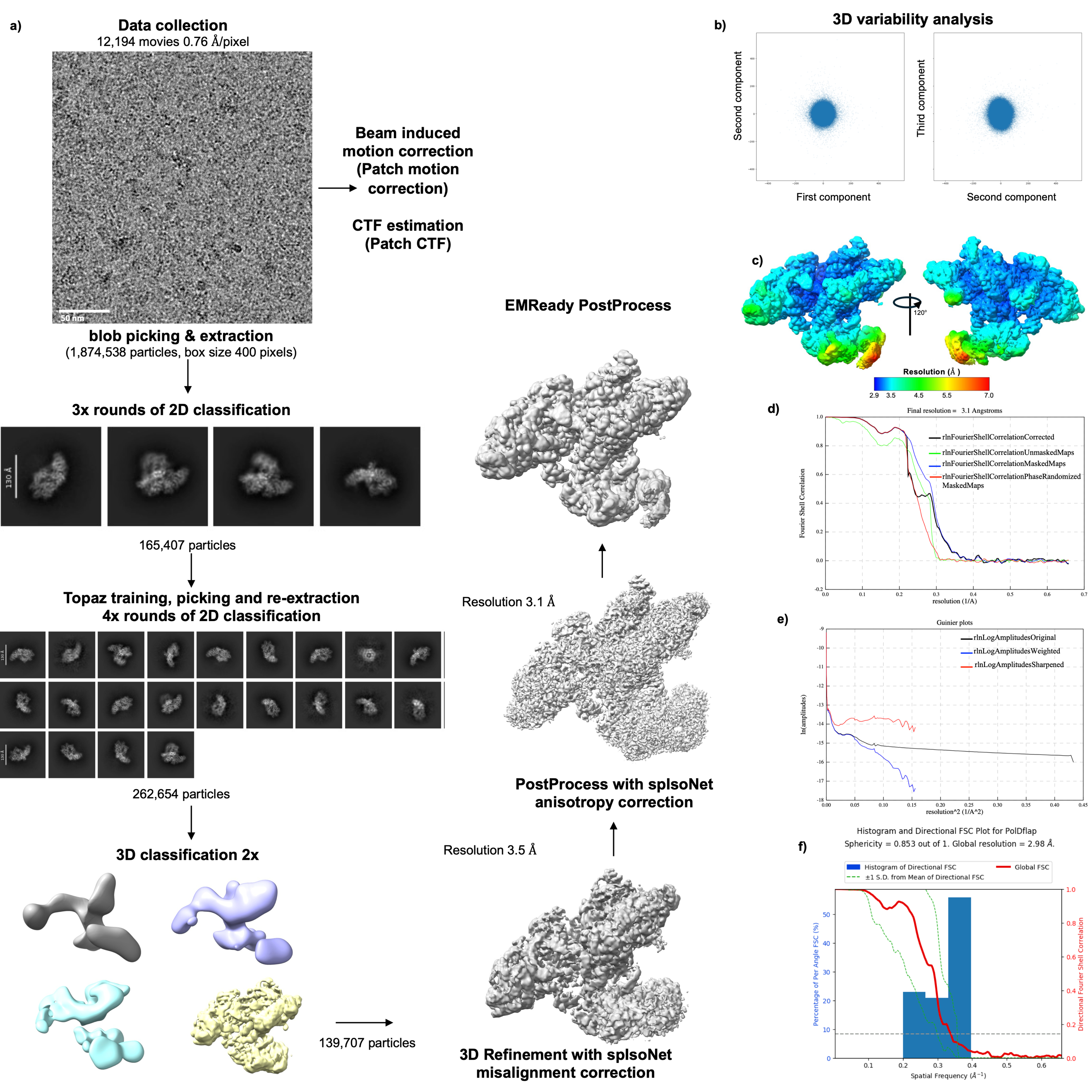

**Extended Data Figure 1**. Single-particle cryo-EM workflow of the strand-displacement Pol δ–PCNA–DNA complex. (a) Data collection and processing workflow. (b) Distribution of particles along the first three components of 3D variability analysis (3DVA), showing a continuous conformational landscape. (c) Local-resolution map of the final reconstruction. (d) Gold-standard Fourier shell correlation (FSC = 0.143). (e) Sharpening/Guinier and map-quality diagnostics from post-processing. (f) Map anisotropy analysis computed by 3DFSC^43^.

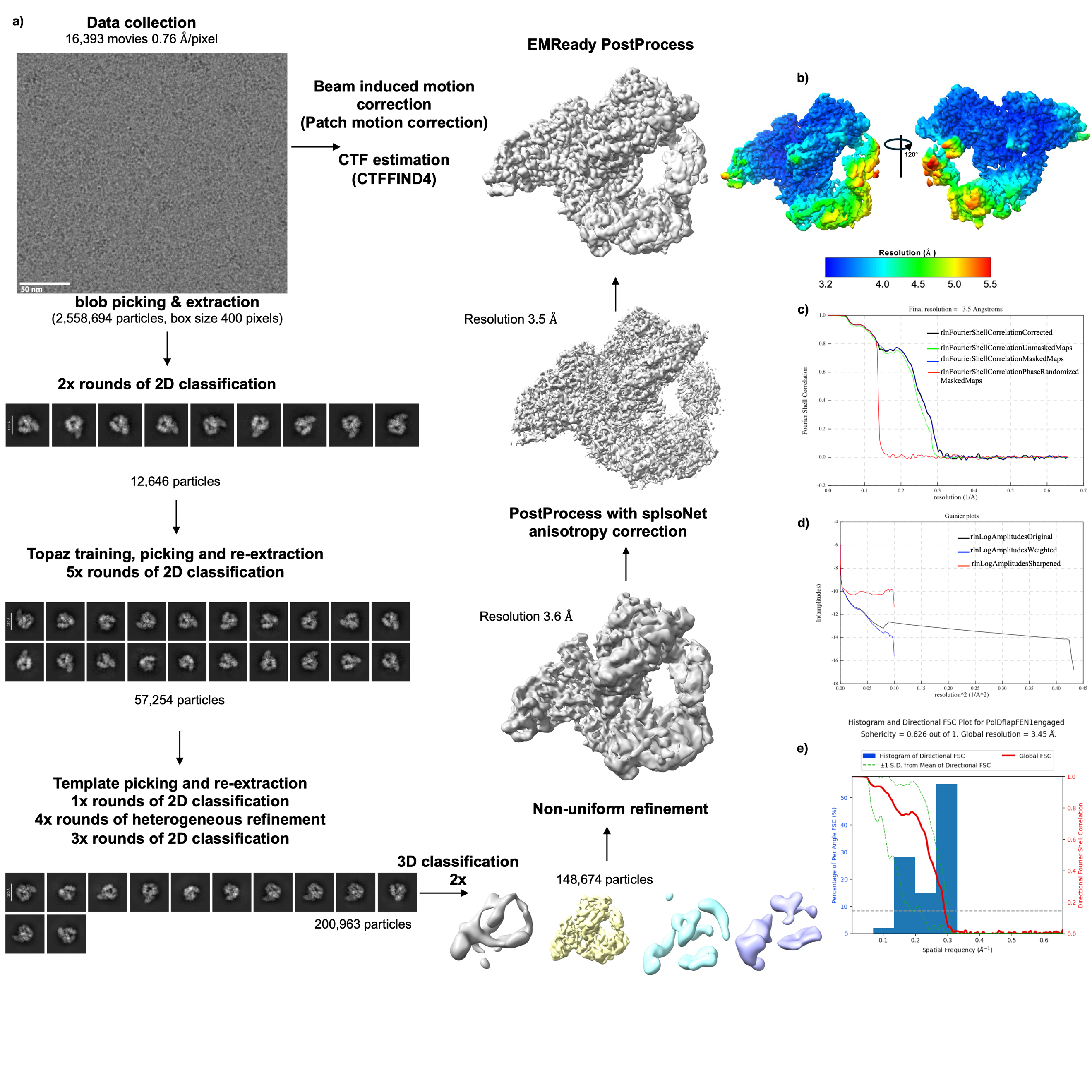

**Extended Data Figure 2**. Single-particle cryo-EM workflow of the Pol δ–DNA–PCNA–FEN–dTTP toolbelt complex. (a) Data collection and processing workflow. (b) Local-resolution map of the final reconstruction. (c) Gold-standard Fourier shell correlation (FSC = 0.143). (d) Sharpening/Guinier and map-quality diagnostics from post-processing. (e) Map anisotropy analysis computed by 3DFSC^43^.

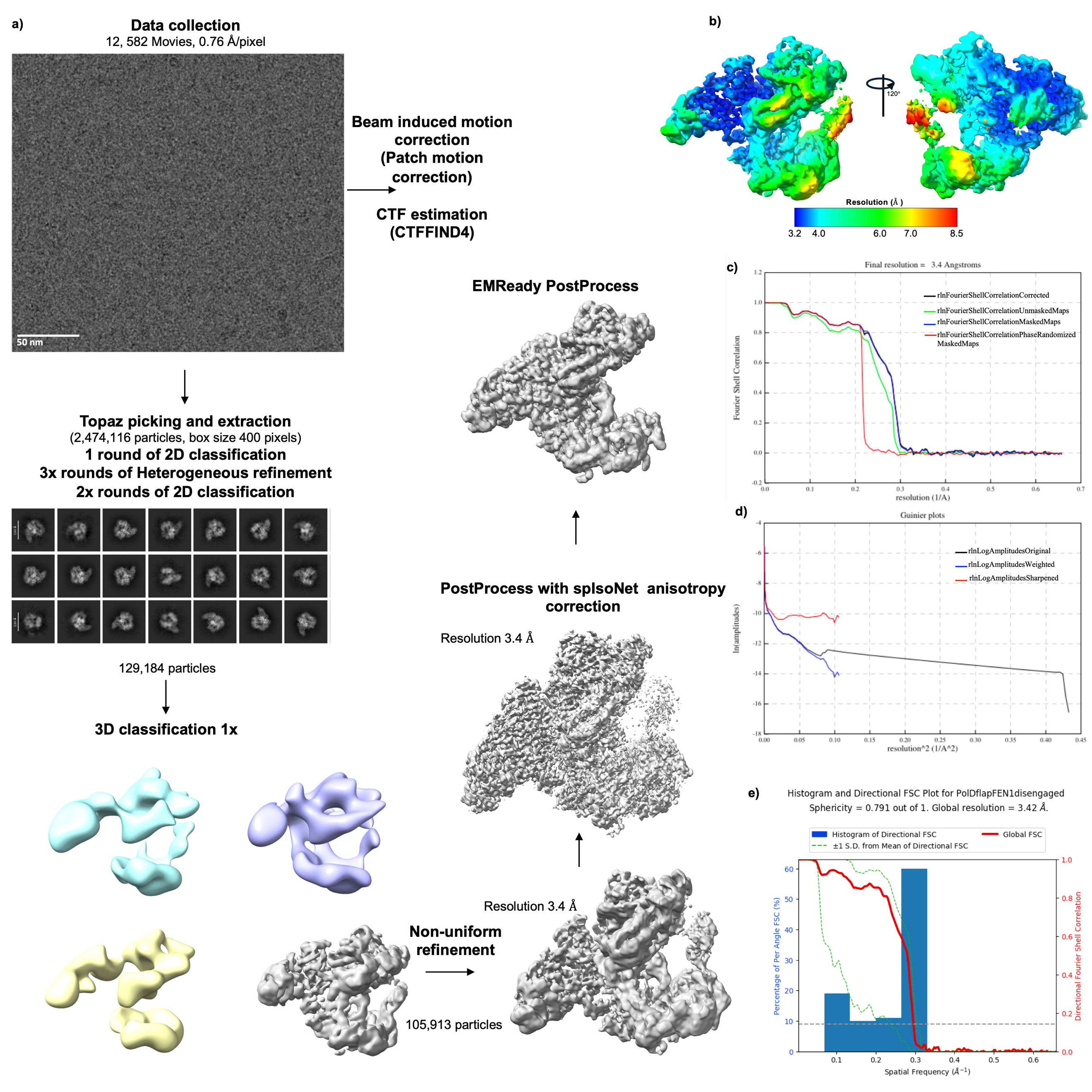

**Extended Data Figure 3**. Single-particle cryo-EM workflow of the Pol δ–DNA–PCNA–FEN1 (no dTTP) toolbelt complex. (a) Data collection and processing workflow. (b) Local-resolution map of the final reconstruction. (c) Gold-standard Fourier shell correlation (FSC = 0.143). (d) Sharpening/Guinier and map-quality diagnostics from post-processing. (e) Map anisotropy analysis computed by 3DFSC^43^.

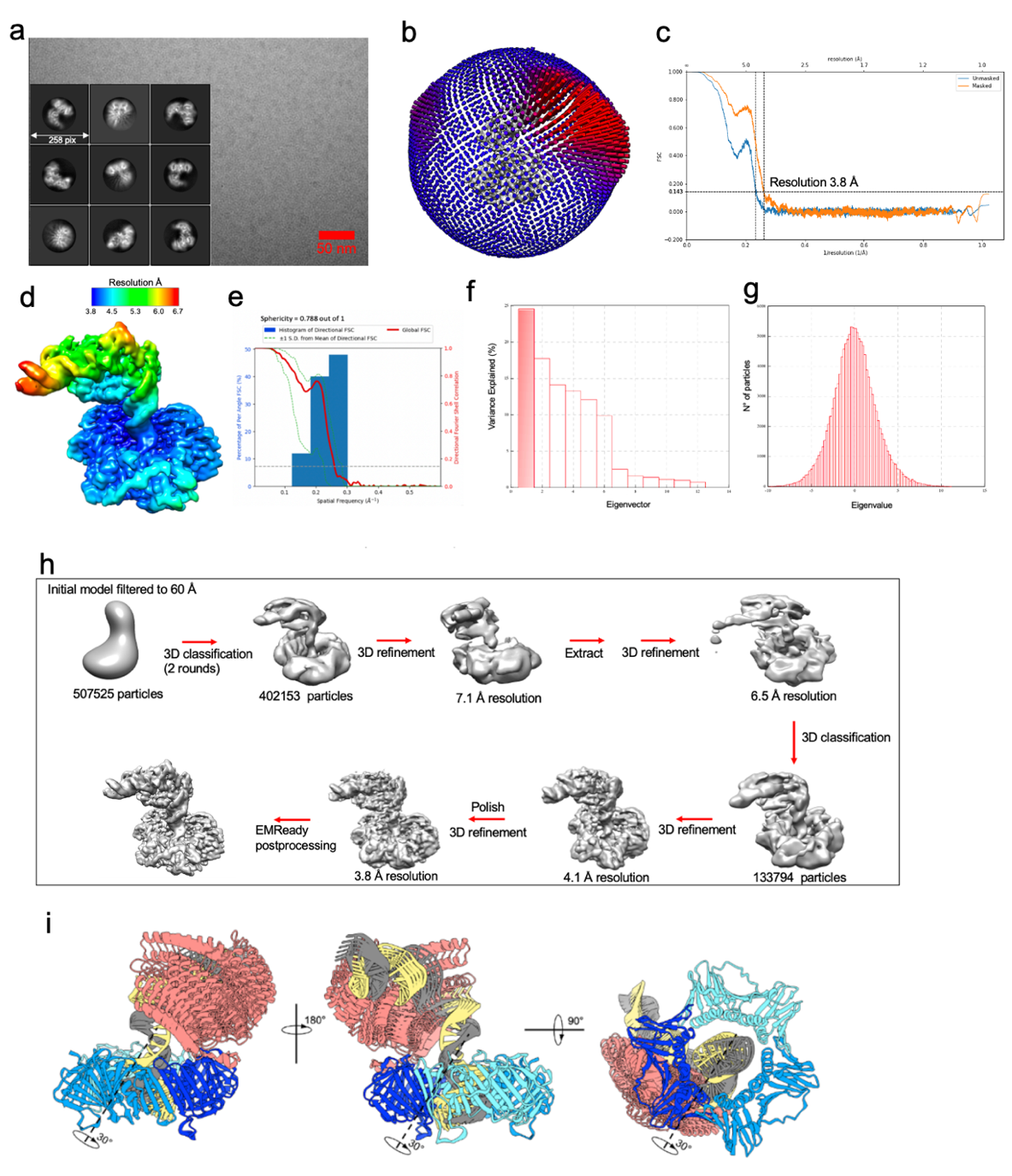

**Extended Data Figure 4. Single-particle cryo-EM analysis of the FEN1–DNA–PCNA complex. (a)** Representative micrograph and selected 2D class averages. **(b)** Angular distribution of particle orientations. **(c)** Gold-standard Fourier shell correlation (FSC = 0.143) indicating a resolution of 3.8 Å. **(d)** Local-resolution map of the final reconstruction. **(e)** Map anisotropy analysis computed by 3DFSC^43^. **(f)** Histogram showing the contribution of eigenvectors to the variance in multibody refinement^44^. **(g)** Histogram of amplitudes along the first eigenvector from multibody refinement, displaying a unimodal distribution indicative of continuous motion. **(h)** Image-processing workflow, including 3D classification, refinement, polishing, and post-processing steps. **(i)** Principal motion derived from the first eigenvector of multibody refinement, corresponding to an ~30° rotation of the FEN1–DNA body relative to PCNA, consistent with flexible tethering via the FEN1 C-terminal PIP box.

**Extended Data Table 1: Cryo-EM data collection, model refinement, and validation statistics**

**Strand displacement dataset**

**3DVA component 000**

Consensus	Frame 1	Frame 20

(EMD-80308)	(EMD-80716)	(EMD-80714)

(PDB 25QR)	(PDB 26KK)	(PDB 26KG)

**Data collection and**

**processing**

Magnification	165,000

Voltage (kV)	300

Total exposure (e-/Å^2^)	41.10

Defocus (µM)	-3.0 to -1.0

Pixel size (Å)	0.76

Symmetry imposed	C1

Initial particles images	2,660,837

Final particles images	139,707

Map resolution (Å)	3.1

FSC threshold	0.143

Map resolution range (Å)	3.0 – 8.0

**Refinement**

Initial model used (PDB code)	6TNY

Model resolution (Å)	3.50	4.00	4.10

FSC threshold	0.5	0.5	0.5

Map sharpening B factor (Å^2^)	-82.2318 (Consensus reconstruction)

Model composition

Non-hydrogen atoms	20101	20101	20102

Protein residues	2399	2399	2399

Ligands	3	3	3

B factor (Å^2^)

Protein	93.19	129.87	132.80

Nucleotide	141.59	221.48	251.46

Ligand	52.59	119.94	112.73

R.m.s deviation

Bond length (Å)	0.007	0.008	0.007

Bond angle (°)	1.302	1.296	1.099

Validation

Molprobity score	1.58	1.64	1.61

Clash score	3.87	4.27	4.40

Poor rotamers (%)	1.07	1.12	0.68

Ramachandran plot (%)

Favored (%)	94.42	94.09	94.13

Allowed (%)	5.58	5.91	5.87

Disallowed (%)	0.00	0.00	0.00

**Extended Data Table 2: Cryo-EM data collection, model refinement, and validation statistics**

**Toolbelt engaged dataset**

(EMD-80285)

(PDB 25PJ)

**Data collection and**

**processing**

Magnification	165,000

Voltage (kV)	300

Total exposure (e-/Å^2^)	43.22

Defocus (µM)	-3.0 to -1.0

Pixel size (Å)	0.76

Symmetry imposed	C1

Initial particles images	6,962,455

Final particles images	148,674

Map resolution (Å)	3.5

FSC threshold	0.143

Map resolution range (Å)	3.3 – 23.3

**Refinement**

Initial model used (PDB code)	6TNY

Model resolution (Å)	3.5

FSC threshold	0.5

Map sharpening B factor (Å^2^)	-169.967

Model composition

Non-hydrogen atoms	22712

Protein residues	2708

Ligands	3

B factor (Å^2^)

Protein	81.70

Nucleotide	125.25

Ligand	59.17

R.m.s deviation

Bond length (Å)	0.009

Bond angle (°)	1.657

Validation

Molprobity score	1.31

Clash score	1.74

Poor rotamers (%)	0.86

Ramachandran plot (%)

Favored (%)	94.50

Allowed (%)	5.50

Disallowed (%)	0.00

**Extended Data Table 3: Cryo-EM data collection, model refinement, and validation statistics**

**Toolbelt disengaged dataset**

(EMD-80304)

(PDB 25QN)

**Data collection and**

**processing**

Magnification	165,000

Voltage (kV)	300

Total exposure (e-/Å^2^)	45.74

Defocus (µM)	-3.0 to -1.0

Pixel size (Å)	0.76

Symmetry imposed	C1

Initial particles images	2,474,116

Final particles images	105,913

Map resolution (Å)	3.4

FSC threshold	0.143

Map resolution range (Å)	3.2 – 14.8

**Refinement**

Initial model used (PDB code)	6TNY

Model resolution (Å)	3.7

FSC threshold	0.5

Map sharpening B factor (Å^2^)	-145.2

Model composition

Non-hydrogen atoms	19621

Protein residues	2394

Ligands	2

B factor (Å^2^)

Protein	102.58

Nucleotide	226.24

Ligand	149.83

R.m.s deviation

Bond length (Å)	0.007

Bond angle (°)	1.290

Validation

Molprobity score	1.47

Clash score	2.81

Poor rotamers (%)	0.87

Ramachandran plot (%)

Favored (%)	94.03

Allowed (%)	5.97

Disallowed (%)	0.00

**Extended Data Table 4: Cryo-EM data collection, model refinement, and validation statistics**

**FEN1-PCNA dataset**

(EMD-80307)

(PDB 25QQ)

**Data collection and**

**processing**

Magnification	105,000

Voltage (kV)	300

Total exposure (e-/Å^2^)	51.00

Defocus (µM)	-2.5 to -1.0

Pixel size (Å)	0.835

Symmetry imposed	C1

Initial particles images	584,121

Final particles images	133,794

Map resolution (Å)	3.9

FSC threshold	0.143

Map resolution range (Å)	3.76 – 6.78

**Refinement**

Initial model used (PDB code)	1UL1, 3Q8K

Model resolution (Å)	3.7

FSC threshold	0.5

Map sharpening B factor (Å^2^)	-131.013

Model composition

Non-hydrogen atoms	10020

Protein residues	1114

Ligands	0

B factor (Å^2^)

Protein	98.95

Nucleotide	133.58

Ligand	-

R.m.s deviation

Bond length (Å)	0.006

Bond angle (°)	1.035

Validation

Molprobity score	1.26

Clash score	1.96

Poor rotamers (%)	0.85

Ramachandran plot (%)

Favored (%)	95.75

Allowed (%)	4.25

Disallowed (%)	0.00
